# Long-distance communication in synthetic bacterial consortia through active signal propagation

**DOI:** 10.1101/321307

**Authors:** James M. Parkin, Richard M. Murray

## Abstract

A synthetic cell-cell signaling circuit should ideally be (1) metabolically lightweight, (2) insulated from endogenous gene networks, and (3) excitable rather than oscillatory or bistable. To accomplish these three features, we propose a synchronized pulse-generating circuit based on the design of published synchronized oscillators. This communication module employs a pulse generator built using Lux-type quorum sensing components and an IFFL transcriptional circuit. Both the input and output of this module are AHLs, the quorum sensing signaling molecule. Cells bearing this module therefore act as an excitable medium, producing a pulse of AHL when stimulated by exogenous AHL. Using simulation and microscopy, we demonstrate how this circuit enables traveling pulses of AHL production through microcolonies growing in two dimensions. Traveling pulses achieve cell-cell communication at longer distances than can be achieved by diffusion of signal from sender to receiver cells and may permit more sophisticated coordination in synthetic consortia.

## 1 Introduction

Quorum sensing circuits, auto-inductive genetic circuits used by wild bacteria for collective decision-making, are a popular platform for designing synthetic communication circuits. Conventionally, quorum sensing circuits estimate cell density. The signaling molecule promotes expression of its associated synthase protein, forming a positive-feedback loop that enables a rapid transition from low to high signal production rate once cell density passes a “quorum” threshold. These natural systems have been applied relatively intact to synthetic applications, such as a circuit that delays expression of an engineered biosynthetic pathway until a quorum has been achieved[1]. With small alterations to the feedback structure, these components can yield complex dynamical behaviors in multi-strain consortia. Two recent examples involve bidirectional signaling between two strains in a co-culture, in one case to achieve sustained oscillations in fluorescent protein expression and in another to recreate the classical predatory-prey dynamical system in an *E. coli* co-culture through AHL-controlled suicide and toxin-rescue circuits [2, 3].

Other applications, however, call for aperiodic and repeated cell-cell communication and cannot rely on the circuits mentioned above. Consider a consortium composed of sensor and actuator microbes embedded in an environment that restricts cell movement, such as intestinal mucus in the mammalian gut. The consortium is intended to persist in this environment and monitor it for a rare chemical event. In response to each event detected by the sensor population, the actuator strain should perform a discrete action, e.g. release a therapeutic small molecule. Without well-mixed conditions, individual sensor cells will differ in exposure to environmental variables of interest and actuators may perceive different concentrations of signal molecule produced by the sensors. These factors reduce the fraction of microbes that participate in signaling, therefore limiting the efficacy of the actuator strains’ impact on the environment.

To overcome the obstacles to group consensus that are inherent to unmixed environments, microbes must collaborate in propagating signals even when they are the signal’s intended recipient. This is because, without sophisticated self-patterning programs, there is no guarantee that all or most of the intended recipient population will receive a signal. Furthermore, participation in the signal propagation must be ephemeral in order to preclude the existence of a steady state with high signal molecule concentration. Signaling via traveling pulses, like action potentials through nervous tissue, achieves these two requirements.

Pulsatile signaling, with active propagation of the pulse by both sensor and actuator strains, would improve efficiency of the consortium described above. Furthermore, pulsatile communication requires only ephemeral investment in the protein components and signal molecules that effect cell-cell signaling, and therefore may be less metabolically demanding for individuals than synchronized bistability. When designing a multi-strain bacterial device or one that may experience a spatially heterogeneous fronts of chemical signals, these features are necessary for reliable, long-distance cell-cell signaling.

Recent work in synchronized transcriptional oscillators has demonstrated that synchronized oscillator strains act as wave generaots when grown in a large enough arena [5, 6]. When syncrhonized oscillator cells were grown in large cell traps, a single cluster in the center of a growing microcolony spontaneously became the focal point for signaling activity of the entire community. This cluster emits, at regular intervals, pulses that travel away from that singular wave source. Taking inspiration from these results, we aim to apply a similar gene circuit to achieve pulsatile communication circuit. This system would enable cells to initiate and transmit communications without permanently altering their internal transcriptional state or spontaneously generating signals at regular intervals.

## 2 Characterizing genetic components in liquid culture

The proposed circuit is an incoherent feedforward loop composed of a Lux-type quorum sensing system augmented with a negative feedforward arm. Lux-type quorum sensing systems consist a transcription activator, an acyl-homoserine lactone (AHL) signal molecule, and an enzyme that synthesizes the signal molecule (synthase). In order to affect transcription, the activator protein must first bind to a signal molecule. The proposed circuit places the signal synthase under the control of a signal-inducible promoter, forming a positive feedback loop on signal molecule concentration. This promoter furthermore contains additional binding sites for a transcriptional repressor. The repressor acting on the promoter is also expressed in response to signal molecule, establishing a negative feedforward arm from AHL concentration to synthase expression mediated via the repressor. This architecture elicits pulsatile synthase expression in response to increases in signal molecule concentration (Fig. 1).

**Figure 1:**
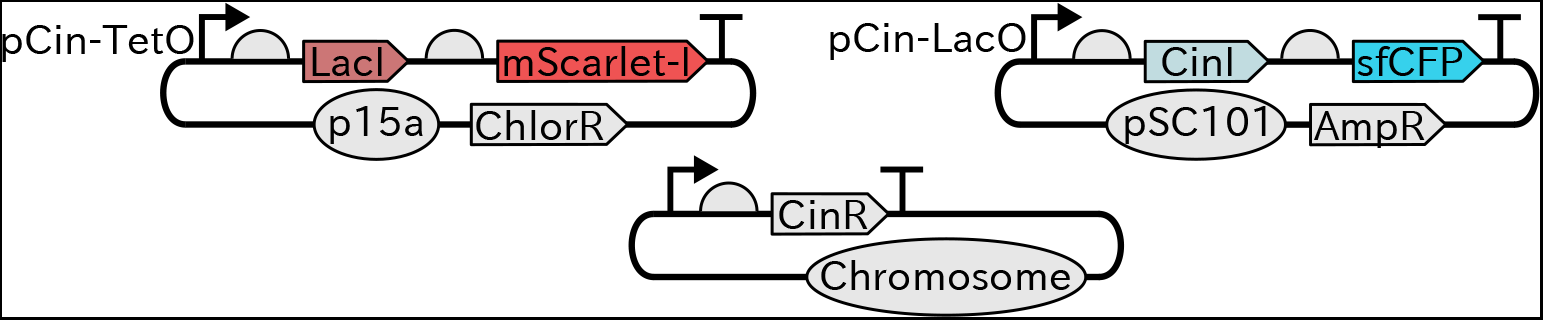
The genetic components of the pulsatile communication circuit. The synthase protein, CinI, produces N-(3-hydroxy-7-*cis*-tetradecanoyl)-L-homoserine lactone (referred to as AHL in the context of the Cin system for convenience) [4]. CinR is expressed constitutively, AHL-bound CinR promotes expression from pCin, and LacI represses CinI transcription when bound to the LacO site. AHL freely diffuses through cell walls.

Plasmids were constructed containing either the synthase or repressor gene. We engineered these plasmids such that fluorescent reporters would be expressed co-transcriptionally with both the synthase and repressor in order to provide a readout of transcription from these two promoters. In all experiments, plasmids were transformed into CY026, an *E. coli* strain that constitutively expresses CinR [2].

We performed liquid culture *in vivo* experiments screening plasmid variants that differed in ribosomal binding site sequences 5’ of the LacI and CinI coding regions. Data from these experiments were fit to a model of the circuit’s behavior. The model assumes Hill-like protein expression as a function of inducer concentration in the media along with cell growth and protein degradation. Two rounds of characterization were performed to optimize circuit behavior. In the first round, strains harboring repressor plasmid candidates were subjected to an induction ladder of the signal chemical. Data from these experiments were fit to a simple model of Hill-like expression as a function of AHL concentration. Because the fluorescent reporter and the repressor genes are bicistronic, the Hill function parameters identified in this process predict the expected expression of the repressor protein, up to the parameter for maximum expression rate. Plasmid candidates that did not fluoresce or that significantly impacted doubling time were removed from further consideration.

In the following round, both repressor and synthase plasmid variants were co-transformed and strains were subjected to a grid of conditions over both AHL and IPTG concentrations, shown in Fig. 2. As expected, synthase expression was higher with increased AHL and IPTG concentrations. It is also clear that both inducers are necessary for significant expression of the synthase-associated fluorescent reporter. This allows for tunable pulse height via IPTG concentration, a useful feature for experiments involving cell-cell signaling. Candidates in this round of characterization were screened for pulsatile expression of the synthase-associated fluorescent reporter and robust cell growth.

**Figure 2:**
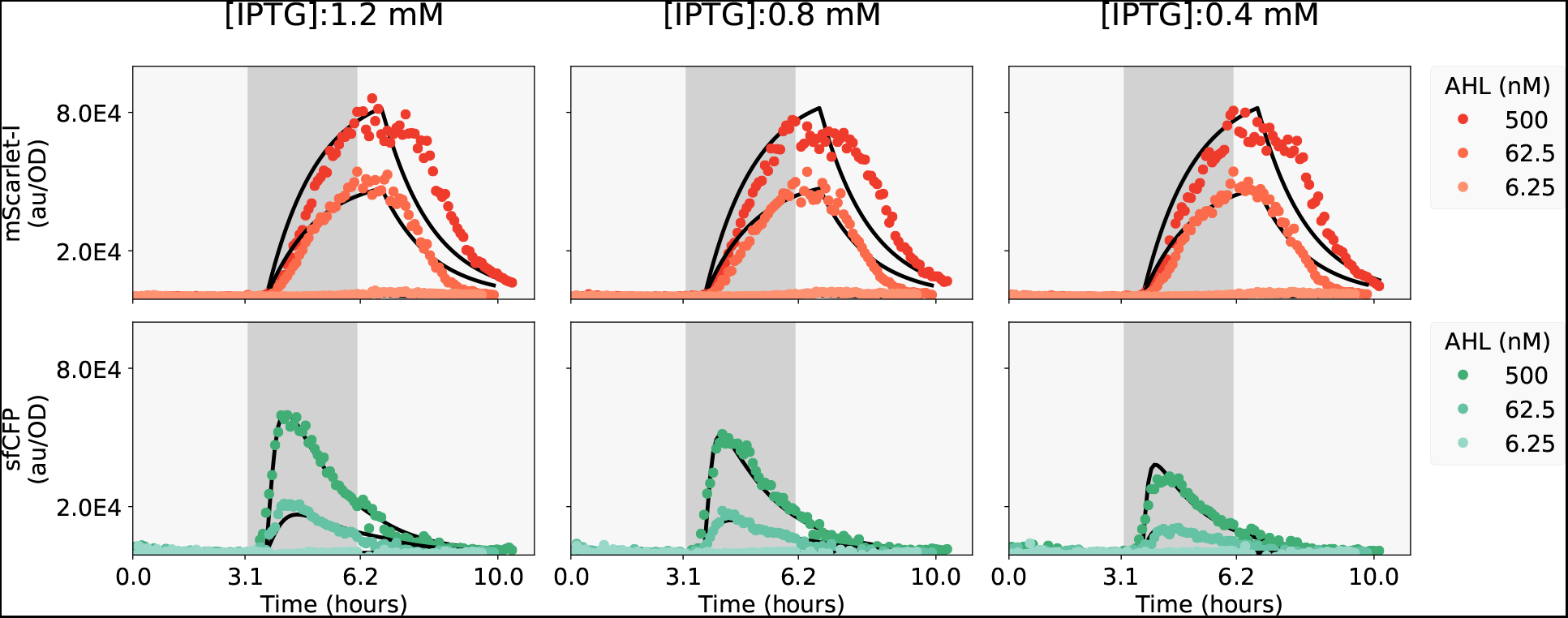
A series of experiments were performed wherein one inducer, IPTG, was kept at a constant concentration while another inducer, AHL, was introduced and then removed. The gray areas indicate the periods when AHL was included in the media. OD-normalized fluorescence data from these experiments, shown in colored points in the above plots, were used to fit parameters by the least-squares method. Data simulated from these fitted model parameters are shown as black lines for comparison. This figure only shows data collected from the candidate strain that demonstrated pulsatile response to AHL and showed the fastest doubling time.

## 3 Modeling circuit behavior in semisolid media and preliminary results

The envisioned use of the pulsatile signaling circuit to support collaboration within synthetic microbial consortia by propagating high-concentration fronts of signal to ensure community consensus in spatially heterogeneous environments. To quantify this capability, we designed a simple experimental scenario to help determine how far a strain can propagate signaling activity from a localized signal source (Fig. 3). Successful propagation should result in transmission velocities at least as great as those observed without active propagation. As shown in Fig. 2, the pulsatile signaling circuit can only produces synthase in the presence of IPTG. This allows for the comparison of transmission with and without active propagation in the same strain.

**Figure 3:**
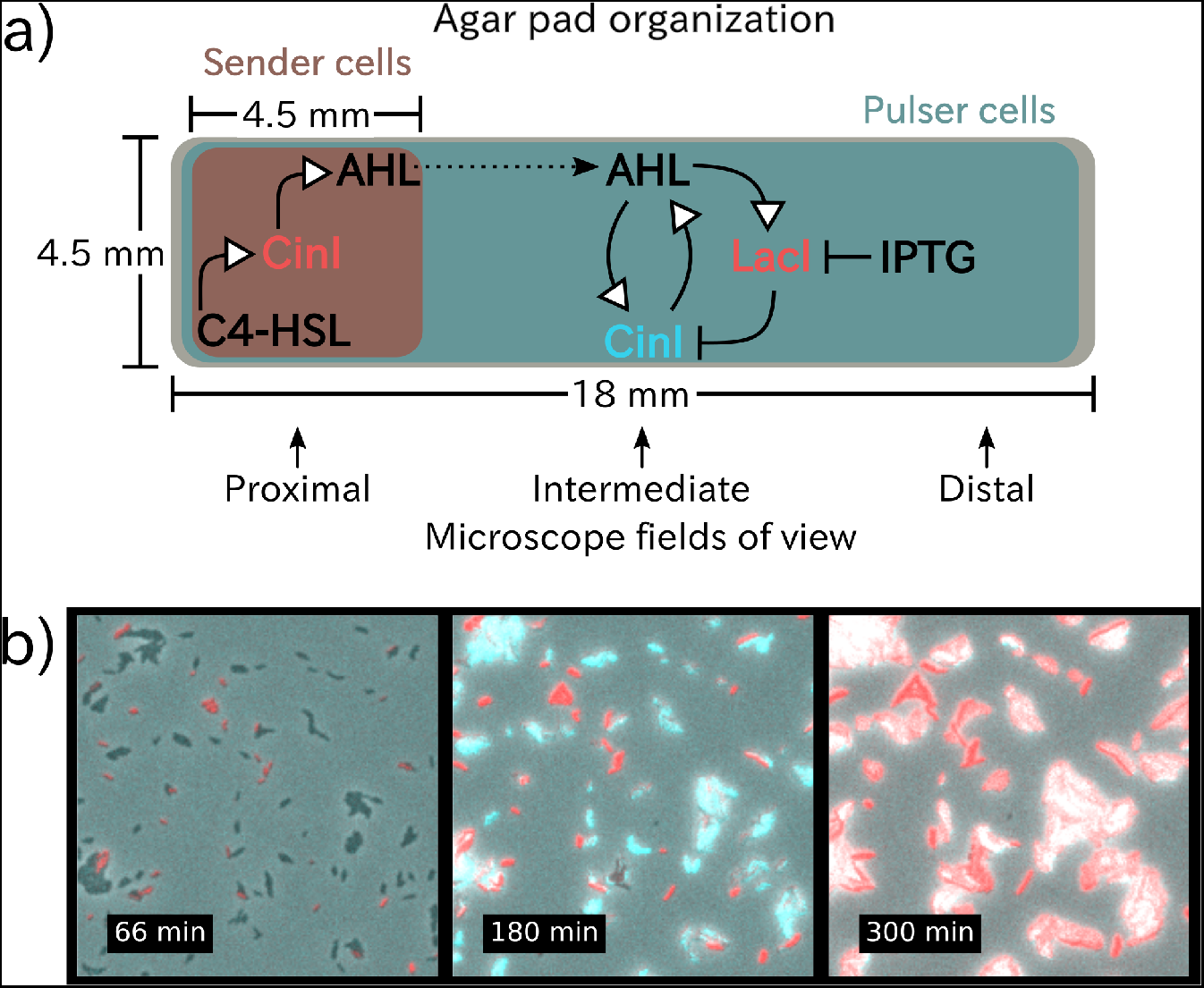
The schematic **a** summarizes the experimental setup described in the text, with red and cyan regions corresponding to the portions of the agar pad occupied by the sender and pulser cell strains, respectively. This schematic also depicts the abstracted transcriptional networks in the sender and pulser strains. Subfigure **b** includes three images from a time-lapse microscopy experiment with C4-HSL and IPTG added to the media, all taken at the proximal position. These images show the order of events in the first phase of cell-cell communication, sender cells releasing AHL that pulser cells detect. At 66 minutes, sender cells are expressing mScarlet-I, indicating that they have been stimulated to produce the synthase protein CinI. The frame at 180 minutes shows neighboring pulser cells expressing sfCFP in response to AHL produced by sender cells. Finally, pulser cells fluoresce in both cyan and red channels at 300 minutes due to expression from both repressor and synthase plasmids, which appears as white in the image.

For both modeling and experimental purposes we apply the organization displayed in Fig. 3 involving two distinct cell strains growing on a rectangular surface of an agar pad. One group of bacteria, the sender cells, are located at one end of a rectangular arena while the pulser cells occupy the entire surface uniformly. If the inducer N-butanoyl-L-homoserine lactone (C4-HSL) is present in the media, sender cells are stimulated to synthesize signal molecule at the beginning of the experiment. AHL then diffuses out of the sender cells and enters neighboring pulser cells. In the case that the agar contains IPTG, they will not only fluoresce but also contribute to the local AHL concentration as well. The result, as shown in simulation (Fig. 4), is a progression of a short band of synthase expression in pulser cells, which emanates from the sender cells and moves towards the far end of the arena.

**Figure 4:**
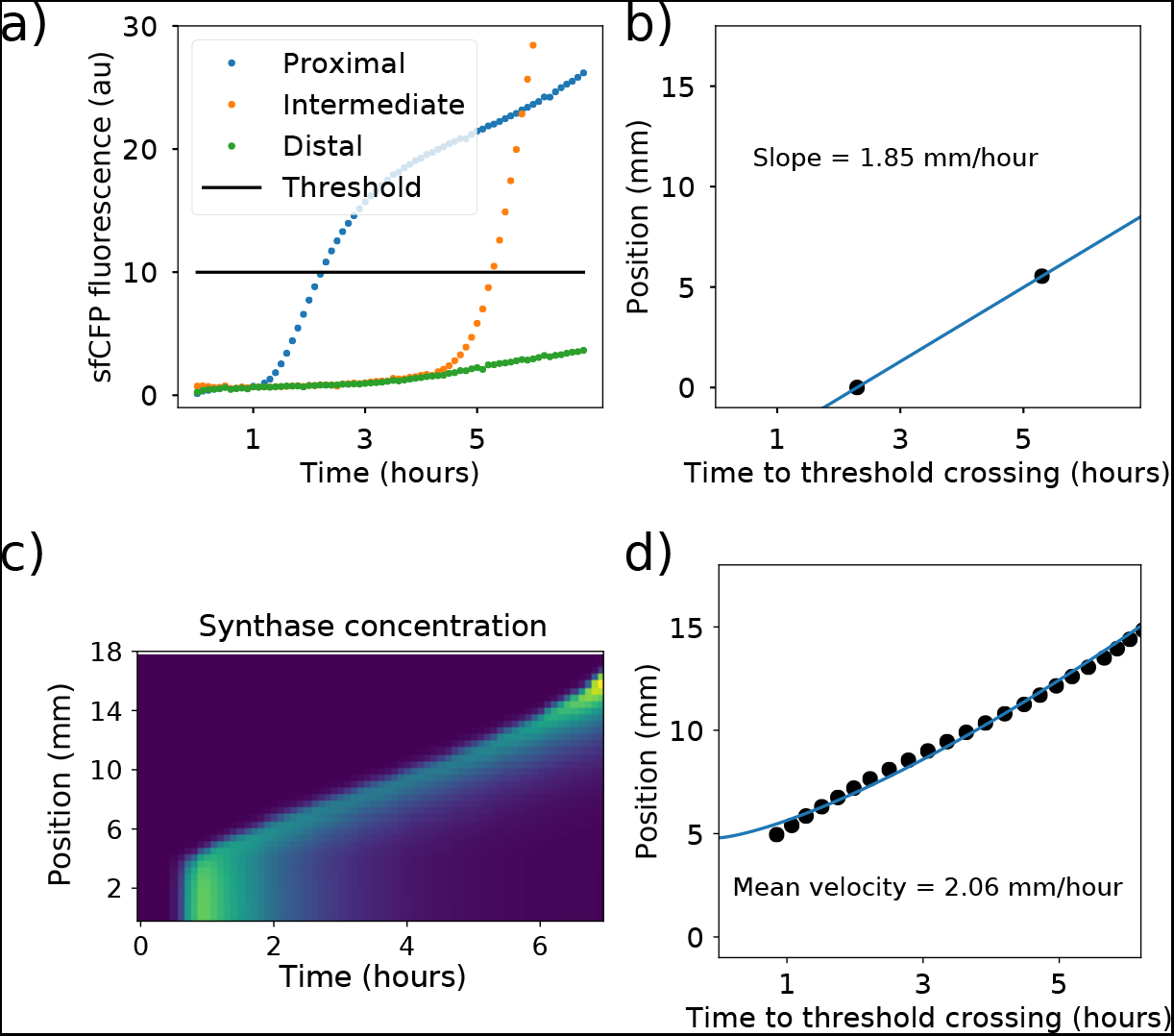
**a** shows the progression of sfCFP fluorescence recorded at three positions of an agar pad organized as in Fig. 3d with activated sender and pulser cells. Fluorescence values were calculated as the mean value over the whole field of view imaged with the corresponding filter set. The threshold value was chosen by hand. **b** shows paired values of threshold crossing times and the estimated distance from the proximal field of view as well as an estimate of the pulse velocity evaluated from the two observed threshold crossing events. Field of view positions are relative to the proximal position. Subplots **c**&**d** depict data from simulation of the intracellular synthase concentration. Simulations treat the agar pad as a 1-dimensional arena. Subfigure **c** shows the relative intracellular synthase concentrations over space and time, with the traveling pulse clearly visible. Threshold crossing times were determined for simulation data and plotted in **d** along with the mean pulse transition velocity. The comparison between simulation and experimental data suggests that the transcriptional parameters identified during liquid culture experiments do not accurately predict behavior of the cells in solid-media experiments. Therefore, transcriptional parameters must be determined through time-lapse microscopy experiments.

**Figure 5:**
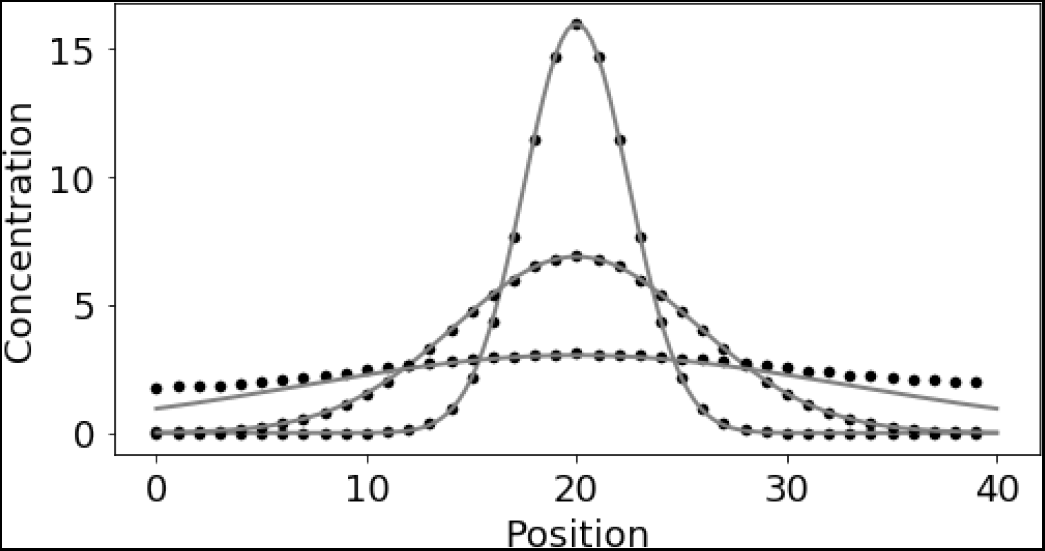
The above plot compares the output of the simulation approach described in this section to the closed-form solution for diffusion of an instantaneous point source of a chemical in one spatial dimension. Both approaches consider the same scenario: an initial concentration of 100 (arbitrary units) at the center of a 40 unit-long arena is allowed to diffuse with a unitary diffusion coefficient. The number of positions considered in the discrete approach is identical to that used when simulating the microscopy experiments (Fig.4). As time increases, the distribution flattens. Each simulated data point falls on top of the closed-form solution, with the exception of the flattest distributions. This is a result of the no-flux boundary conditions imposed in the simulation. Because the closed-form solution was derived assuming an unbounded domain, the concentration will continue to diminish and gradually approach zero. In the simulation, however, the concentration at each position will approach a nonzero steady state.

We adapted a model described previously to the experiments described here [7]. This model was developed to describe the growth of colonies on agar plates as well as communication between colonies via production and diffusion of quorum sensing molecules. Parameters describing cell growth and AHL diffusion were taken directly from this model, as it has been previously verified on the same molecules and *E. coli* strain to be used in this work. The transcriptional parameters, on the other hand, were derived from the liquid culture experiments described above. Applying these parameters to the model produced in simulation the traveling pulse behavior we expected, suggesting that the genetic components optimized through liquid culture experiments would show the desired behavior when applied to 2D environments.

To perform *in vivo* experiments, sender cells were cloned such that they produce the fluorescent protein mScarlet-I and AHL synthase CinI in response to C4-HSL, another quorum sensing signaling molecule. Pulser cells were identical to those used in liquid culture characterization experiments. In each microscopy experiment, cells were deposited on the agar pad in the same pattern and from the same source cultures. Composition of the agar pads varied only in the presence or absence of IPTG and C4-HSL. IPTG is necessary for CinI expression from pulser cells and C4-HSL is necessary for CinI expression from sender cells.

Preliminary data collected from time-lapse microscopy of cells organized on an agar pad according to the above description suggests that the circuit characterized through liquid culture experiments successfully increased the signaling distance of sender cells. Three experimental conditions were assayed to determine if active propagation would extend the signaling distance of the sender cells: +IPTG –C4-HSL, +IPTG +C4-HSL, –IPTG +C4-HSL. Under the first set of conditions, + IPTG – C4-HSL, the sender cells are inactive, so neither the sender nor pulser cells should fluoresce. The second set of conditions should elicit the traveling pulse behavior from proximal to distal ends of the agar pad. The third condition set supports a simple sender-receiver relationship in which the pulser cells create LacI and mScarlet-I in response to AHL, but not CinI nor sfCFP. This case would demonstrate the signaling distance achievable through simple diffusion of AHL from the proximal end of the pad.

Figure 4 shows fluorescence data collected from cells at the proximal, intermediate, and distal ends of the agar pad relative to the position of the sender cells. By the end of 10 hours, only the second condition lead to significant fluorescence from cells in the intermediate position. This indicates that the signaling distance achieved in the second condition was longer than in the third condition, suggesting that active propagation did significantly increase the transmission distance.

Note that the velocity of the traveling pulse observed from time-lapse microscopy was estimated to be 1.85 mm/hr, while the velocity demonstrated in simulation using parameters derived from liquid culture characterization predicted a velocity of 2.06 mm/hr. This suggest that, while liquid culture characterization was necessary to determine appropriate genetic material for pulse generation, a second set of characterization experiments performed using single-cell microscopy will be necessary to find model parameters that are predictive of the spatial signaling behavior on agar pads.

## 4 Methods

Liquid culture experiments were performed in a Biotek Synergy H1F plate reader using M9CA minimal media (Teknova product code M8010–06) with 100 *μ*g/mL ampicillin and 34 *μ*g/mL chloramphenicol. Starter cultures were inoculated into M9CA media from single colonies picked from an agar plate. Inoculated cultures were shaked and incubated at 37°C until the OD600 reached 0.3. The experimental wells were then prepared by diluting starter cultures 1:20 into a final volume of 500*μ*L in a 96-well glass-bottom plate. When necessary, inducer chemicals were added to wells using an Echo 525 acoustic liquid handler before the addition of cell culture and broth.

At two points during the liquid culture experiments, inducer concentrations were altered by diluting and washing experimental cell cultures. Washes were performed by first pelleting the full 500*μ*L of culture from an experimental well, discarding the supernatant, and then resuspending in 15mL of PBS. These wash-discard-resuspend cycles were repeated twice more for a total of three wash cycles. 500 *μ*L of M9CA broth was used in the final resuspension step to return the culture to approximately the same density as before the wash steps. Subsequent experimental wells were prepared by first depositing inducer chemicals as needed with the Echo 525 liquid handler, followed by cell culture and sterile M9CA broth in a 1:20 ratio.

Analysis of data collected from plate reader experiments was performed using custom Python scripts. Background fluorescence and OD600 were determined by measuring these quantities in wells prepared without inducer chemicals, which previous experiments had shown to be identical in fluorescence to cells without fluorescent reporters. These background signals were determined to be time-varying, resulting from either background fluorescence from growing cells or from broth exposed to air. OD600-normalized fluorescence values (fluorescence divided by OD600) were calculated using background-subtracted data.

Microscopy experiments were performed using agar pads prepared according to the protocol described in [8]. Pads were prepared with inducers introduced to the molten agar and without antibiotics. When added to molten agar, final IPTG concentration was 1 mM and final C4-HSL concentration was 10 *μ*M. Images were acquired using an Olympus IX81 inverted microscope through a UPlanFl10XPh objective and Chroma filters 41027 and 310442V2 for mScarlet-I and sfCFP, respectively. Sample temperature was held at 29°C for the duration of the time-lapse microscopy.

Analysis of data collected from microscopy experiments was performed using custom Python scripts. As mentioned in the caption to Fig. 4, fluorescence values were calculated as the mean over the entire field of view at each time point, position, and filter set. Fluorescence values from the distal position of the inactive sender and receiver cell agar pad were taken as the background signal at each time point for all positions and conditions. Data presented in 4 are background-subtracted.

## 5 Model equations and parameters

### Liquid culture model

We here describe the ordinary differential equations model for pulse-generator circuit behavior in liquid culture. The model species are the cellular concentrations of synthase and repressor proteins, in units of OD-normalized fluorescence (referred to as arbitrary units or a.u.). Two terms make up the derivative expressions for the two protein species: production and degradation.

We assume Hill-like protein production and constant proportional degradation. Repressor expression is activated by AHL, therefore the production term is an increasing Hill function. 
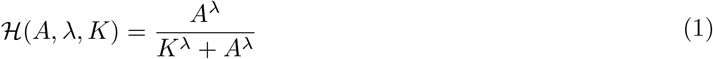
Synthase expression is activated by AHL and repressed by the repressor protein. We use a decreasing Hill function to model the dependence of synthase expression on repressor protein concentration. 
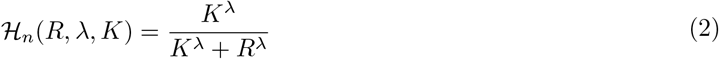

The effective repressor concentration, however, is modulated by IPTG concentration. IPTG binds to LacI, preventing it from acting on operator regions in promoter sequences. To model the effect of IPTG, we calculate the effective repressor concentration via a decreasing Hill function. 
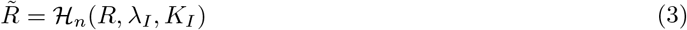

The full set of differential equations is as follows: 
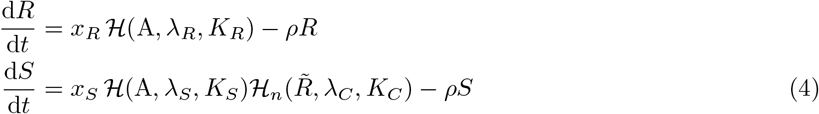

With expression terms *x*, degradation rates *ρ*, Hill coefficients *λ*, and critical concentrations *K*. While the synthase will contribute to the concentration of AHL, we do not model this effect. The amount of time that cells are expressing synthase is small enough that the AHL produced by cells is of little consequence to the trajectories of sfCFP or mScarlet-I fluorescence. Model parameters determined by least-squares fitting to the data presented in Fig. 2 are shown below.

| Parameter | Value | Units |
| --- | --- | --- |
| $x_R$ | 18.6 | a.u. seconds<sup>-1</sup> |
| $x_S$ | 64.3 | a.u. seconds<sup>-1</sup> |
| $\lambda_C$ | 3.27 | |
| $\lambda_R$ | 2.30 | |
| $\lambda_S$ | 4.98 | |
| $K_C$ | 691 | a.u. |
| $K_R$ | 55.3 | nM |
| $K_S$ | 85.9 | nM |
| $\lambda_I$ | 0.499 | |
| $K_I$ | 1.72 | μM |
| $\rho$ | 2.02E-4 | seconds<sup>-1</sup> |
| delay | 2010 | seconds |

### Solid media model

Building on the model described in the previous section, we constructed a reaction-diffusion model to describe the behavior of cells growing on agar. This simulation approach was adapted from one used in a related project [7]. The model considers nutrient-dependent cell growth, nutrient-dependent protein production, signal synthesis via synthase proteins, signal diffusion, cell diffusion, and nutrient diffusion. As the circuit is relatively simple, we elected for the model to use a discretized approximation of a continuous reaction-diffusion system. In this discretization, the agar plate is compartmentalized into square chambers 450 *μ*m to a side. The concentration of model species takes on a single value within each voxel. To calculate the discretized approximation of diffusion, the model calculates the rate of change in molecule concentration by applying Ficks’ first law to the surfaces dividing adjacent rectangular chambers. For indices (i, j) and (x, y) on a grid defining chamber locations on a plate P, and scale term *d_x_*, discrete approximation of the spatial derivative of a species*s* takes the form (ignoring the diffusion coefficient): 
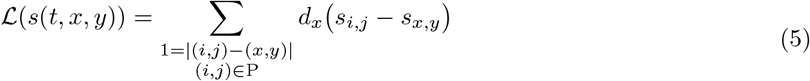

The scale term *d_x_* is derived from the discrete approximation of Fick’s first law. Consider *s*, the concentration of a diffusible species in a cubic chamber with edge length *w* and diffusion coefficient *D*. The change in concentration due to diffusion through one chamber face can be calculated first by accounting for the molecular flux and then determining the resulting change in concentration. Note that here, the center-to-center spacing of the chambers *x* is the same as the chamber edge length *w*. 
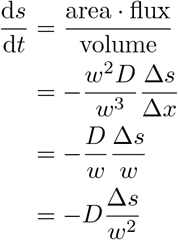

The term *d_x_* used in Equation 5 is equal to 1 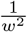, where *w* is the chamber edge length. Furthermore, the simulation includes no-flux boundary conditions 5. All other model equations make use of Hill functions to model nonlinear, bounded activity promoted by small molecules and modulated by the activity of transcriptional repressors (Equations 1&2).

Cell growth, protein production, and signal molecule production all depend on nutrient availability. Each time derivative of these species involves production terms that are proportional to a production parameter, the nutrient availability in that chamber, and the concentration of the producing species (e.g. synthase is the producing species for AHL). Nutrient availability is a nonlinear function of the local nutrient concentration, defined by a Hill function. 
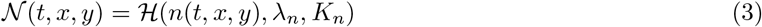

While protein production and cell growth depend on nutrient availability, only cell growth actually consumes nutrients. This feature is anchored in the assumption that the metabolic investment in the circuit’s component proteins is minor compared to all other cellular activity. Protein dilution and degradation are separate terms in the reaction-diffusion model, whereas they were one term in the liquid culture model.

The simulated agar pad is slightly different from the pad described in Fig. 3. In simulation, the cell density is homogeneous. Therefore, simulating the 4.5mm × 18mm agar pad would result in wasted computation, as diffusion along the shorter pad dimension would always be zero. To reduce unnecessary computation, the simulated agar pad is only two chambers, or 0.9 mm, wide.

Two *E. coli* strains are deployed in the solid-media experiments. Experimentally, the sender cells are deposited on top of a portion of the uniformly-disseminated pulser cells. For modeling simplicity, the two strains occupy different regions of the arena. In simulation, the agar pad is divided along a center line running from the proximal to distal end, with sender cells on one side of the line and pulser cells on the other. Each strip is one chamber wide and forty chambers long. The sender cells only occupy the proximal quarter of their alloted strip while the pulser cells occupy the entire length of theirs. This simplification reduces unnecessary computation, as noted above, while ensuring rapid diffusion of nutrient and AHL from sender to pulser cells.

Protein species in the model represent intracellular concentrations. To codify how the two strains contribute to local protein production, the model applies a threshold to the local strain densities. 
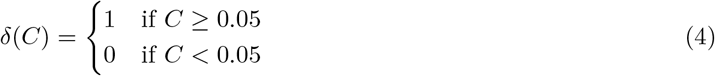

Pulser cells, for example, only produce synthase in the presence of AHL and an absence of repressor. Sender cells, on the other hand, produce synthase constitutively. Each production term (Hill-like or constant) is multiplied by the threshold function evaluated on the density of the associated strain to determine which production behavior to engage at a given location.

The full system of ordinary differential equations defining the model is listed here. Furthermore, the model is built with realistic parameters and units for AHL diffusion and concentration, including time, but we leave the units describing cell density, protein quantity, and nutrient concentration arbitrary. 
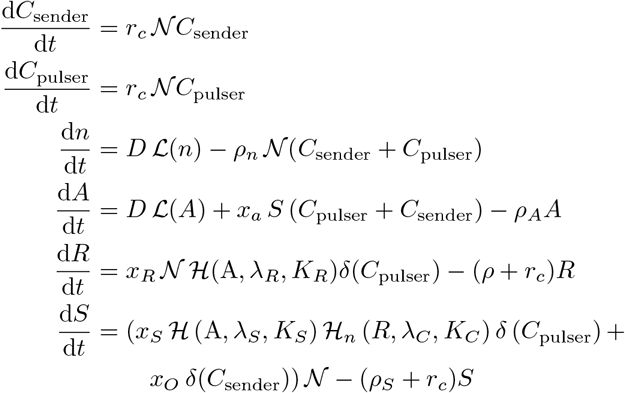

The parameter values used to generate the simulation data shown in Fig. 4 are shown in the table below. The term od0 refers to the initial cell density, in units of optical density as measured.

| Parameter | Value | Units |
| --- | --- | --- |
| $d_x$ | 4.93 | $\text{mm}^{-2}$ |
| $r_c$ | 0.008 | $\text{min}^{-1}$ |
| $\rho$ | 0.008 | $\text{min}^{-1}$ |
| $K_n$ | 15 | |
| $D_n$ | 0.03 | $\text{mm}^2 \text{ min}^{-1}$ |
| $\rho_n$ | 1 | $\text{min}^{-1}$ |
| $D_A$ | 0.03 | $\text{mm}^2 \text{ min}^{-1}$ |
| $x_A$ | 0.08 | $\text{nM min}^{-1} \text{ OD}^{-1} \text{ a.u.}^{-1}$ |
| $x_S$ | 500 | $\text{a.u. min}^{-1}$ |
| $x_O$ | 200 | $\text{a.u. min}^{-1}$ |
| $x_R$ | 1200 | $\text{a.u. min}^{-1}$ |
| $\lambda_S$ | 3.2 | |
| $K_S$ | 100 | $\text{nM}$ |
| $\lambda_R$ | 2.3 | |
| $K_R$ | 55 | $\text{nM}$ |
| $\lambda_C$ | 3.2 | |
| $K_C$ | 2700 | $\text{a.u.}$ |
| $\rho_A$ | 0.005 | $\text{min}^{-1}$ |
| od0 | 0.1 | $\text{OD}$ |

## 6 Acknowledgments

James M. Parkin is supported by the Institute for Collaborative Biotechnologies through grant W911NF-09–0001 from the U.S. Army Research Office. The content of the information does not necessarily reflect the position or the policy of the Government, and no official endorsement should be inferred. Plasmid vectors and non-coding regions were provided as a generous gift of Douglas Densmore at the Cross-disciplinary Integration of Design Automation Research lab (Addgene Kit # 1000000059). Quorum sensing promoters, quorum sensing protein coding sequences, and strain CY026 were provided as a generous gift from Matthew Bennet (Addgene Plasmid #65954, #65952, Bacterial Strain #72340).

